# NR4A3 is oxidative stress responsible transcription factor through HMOX1, and also controls cell cycle through CCNE1 and CDK2 in pancreatic islet derived 1.1B4 cells

**DOI:** 10.1101/2021.08.20.457070

**Authors:** Motoko Nakayama, Etsuko Ueta, Mitsuru Yoshida, Yuri Shimizu, Reiko Oguchi, Atsuko Tokuda, Yasuko Sone, Yuri Nomi, Yuzuru Otsuka

## Abstract

The mechanism of antioxidant defense system is still controversial. As islet ß-cell is weak in oxidative condition, that causes diabetes mellitus, therefore, antioxidant defense system of human pancreatic islet derived 1.1B4 cell was analyzed. Cells were exposed to H_2_O_2_ and comprehensive gene expression was analyzed by Agilent human microarray. *HMOX1* and *NR4A3*, member of orphan receptor, were up-regulated. Therefore, *NR4A3* was knocked down with siRNA, then analyzed gene expression by microarray, and found that the knocked down cells were weak in oxidative stress. *HMOX1* expression was strongly inhibited by siRNA of *NR4A3*, and *NR4A3* responsible sequence of aaggtca was found near the *HMOX1* gene, suggesting *NR4A3* is oxidative stress responsible transcription factor through *HMOX1* expression. The expression of *CCNE1* and *CDK2* was also inhibited by knocked down of *NR4A3*, it is suggested *NR4A3* is also important transcription factor for cell growth regulation.

**Graphical abstract:** 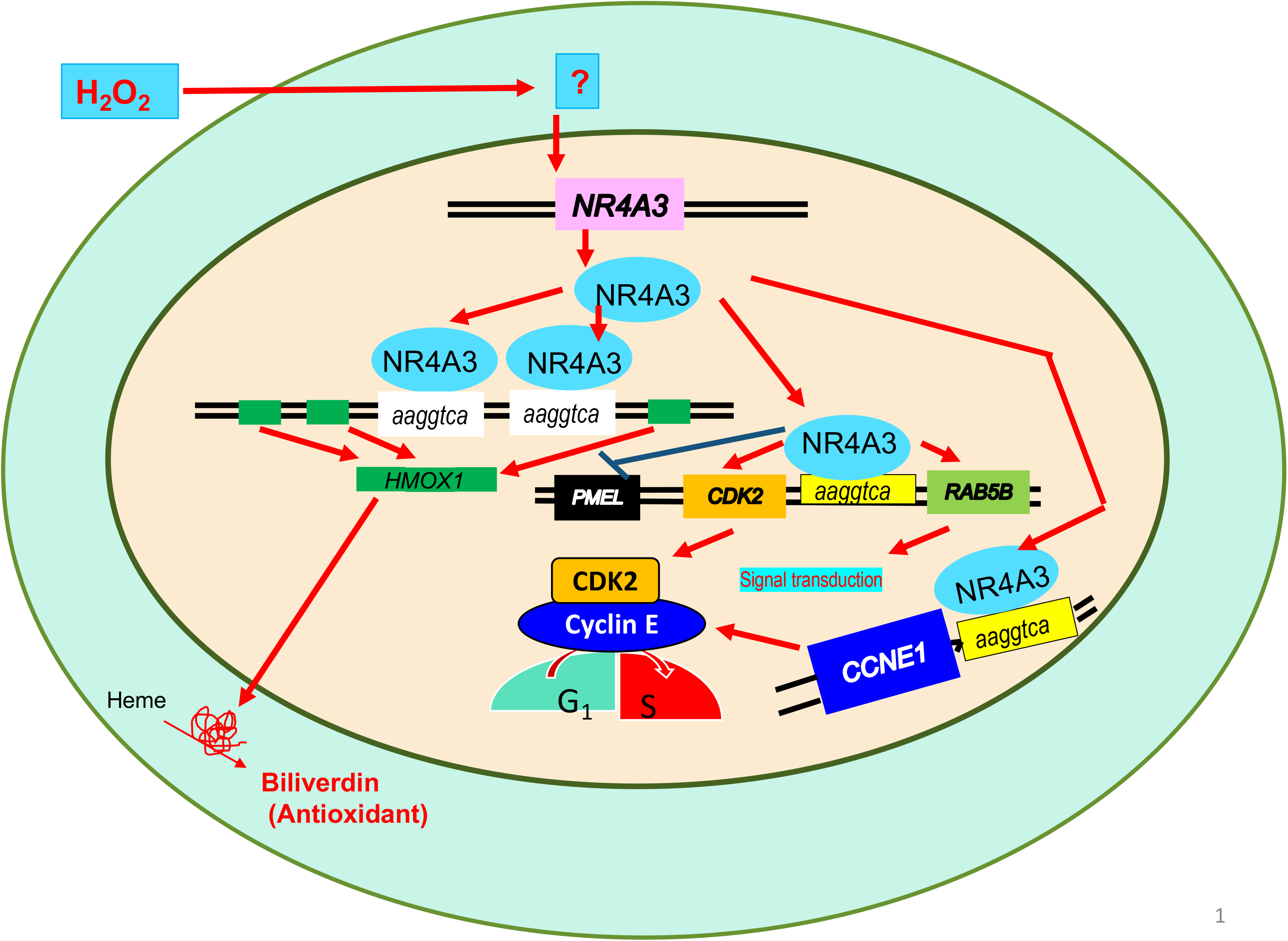
Hydrogen peroxide induces NR4A3 and binds to aaggtca sequence of HMOX1, and increased transcription of HMOX1. Resulting heme oxygenase produces biliverdin, antioxidants, from heme. NR4A3 also bind to aaggtca sequence of CDK2 and CCNE1, resulting CDK2 and Cyclin E. CDK2 bind to cyclin E and cell goes from G1 to S phase.

## INTRODUCTION

The relentless decline in ß-cell function frequently observed in type 2 diabetic patients and loss of functional ß-cell mass is a hallmark of type 1 and type 2 diabetes, and methods for restoring these cells are needed (Tessem et al. 2014). However, despite optimal drug management, loss of ß-cell is not possible to stop, and has variously been attributed to glucose toxicity and lipo-toxicity. The former theory posits hyperglycemia, that elevated glucose concentrations increase levels of reactive oxygen species (ROS) in ß-cell (Robertson 2004), which takes place within multiple mitochondrial and non-mitochondrial pathways. For example, high concentrations of glucose in vitro increase intracellular peroxide levels in islets, and decrease insulin expression by decreasing *PDX-1* and *MafA* expression (Robertson and Harmon 2006). In their paper, six biochemical pathways along which glucose metabolism suggested to form ROS. One of pathway is glycation that is hyperglycemia produces glycation end product (AGE) that increases ROS, as AGE-BSA is known to increase production of ROS, and its apoptogenic effect was blocked by the antioxidant N-acetylcysteine (Weinberg et al. 2014). In physiologic concentrations, endogenous ROS help to maintain homeostasis, however, when ROS accumulate in excess for prolonged periods of time, they cause chronic oxidative stress and adverse effects. Thus, increased glucose produces ROS, and pancreatic ß-cells are more sensitive to oxidative stress than other cell types because the expressions of antioxidant enzymes are low levels in pancreatic ß-cell (Robertson 2004). Therefore, oxidative stress leads cellular damage to ß-cell dysfunction. The effect of oxidative stress, however, on the ß-cell is not understood well. Nitric oxide, peroxynitrite, hydrogen peroxide and other oxygen-reactive species might be involved in ß-cell destruction during diabetes development (Mathis et al. 2001). Oxidative stress induces gene expression in heme oxygenase 1 (HMOX1), and the HMOX1 gene (*HMOX1*) is frequently activated under a variety of cellular stress conditions, by four pathways including heat-shock factor, nuclear factor-kappa B, nuclear factor E2-related factor 2 (NRF2), and activator protein-1 families, are arguably the most important regulators of the cellular stress response in vertebrates (Alam and Cook 2007). One of them, the Kelch-Like ECH-Associated Protein 1 (KEAP1)-NRF2 pathway in which pathway, NRF2 transcription factor directly bind to antioxidant responsible element (ARE) to induce antioxidant enzymes (Nguyen et al. 2009). However, another unknown pathway may be involved. For example, chronically excessive glucose and ROS levels can cause decreased insulin gene expression via loss of the transcription factors pancreatic and duodenal homeobox 1 (PDX1) and v-maf avian musculoaponeurotic fibrosarcoma oncogene homolog A (MAFA) and can also accelerate rates of apoptosis (Robertson 2004).

High glucose and oxidative stress induce cell damage not only to ß-cell but also to other cells. There are many transcription factors and sensing proteins. Oxidants such as H_2_O_2_ can damage proteins, regulate transcription factors, and sometimes induces cell apoptosis (Marinho et al. 2014). In his review, the regulatory mechanisms by which H2O2 modulates the activity of transcription factors in bacteria (*OxyR* and *PerR*), lower eukaryotes (*Yap1, Maf1, Hsf1* and *Msn2/4*) and mammalian cells (*AP-1, NRF2, CREB, HSF1, HIF-1, TP53, NF-κB, NOTCH, SP1* and *SCREB-1*) are summarized. Furthermore, Klamt *et al*. (Klamt et al. 2009) found that the actin-binding protein cofilin is a key target of oxidation. When oxidation of this single regulatory protein is prevented, oxidant-induced apoptosis is inhibited.

As described above, high glucose increases ROS, and antioxidants repairs ß-cells undergoing damage by oxidative stress, however, molecular mechanism of ROS on the islet ß-cell is not well understood. Therefore, in this paper we focus on the mechanism of gene expression by ROS using H_2_O_2_ in human pancreatic ß-cells derived hybrid cell. Because, transformed cell used for in vitro experiment, the cells sometime lost their characteristic especially apoptosis pathway, otherwise cell will die. Furthermore, human normal islet cells are not available easily, 1.1B4 cells, the hybrid cell of normal islet cell with ß-cell derived transformed PANC-1 cell was used here (McCluskey et al. 2011). In this paper we analyzed the effect of H_2_O_2_ on the gene expression of 1.1B4 with Agilent microarray. Then key protein was knock down by siRNA method and analyzed cell viability under the H_2_O_2_ addition. And gene expression of siRNA treated cells was analyzed by microarray again to find how this knock down of the key protein effect gene expression.

## RESULTS

### Effect of H_2_O_2_ on 1.1B4 cells growth and gene expression analyzed by DNA microarray

Human pancreatic islet derived 1.1B4 cells were exposed to 10, 20, 30, 40, 50, 100, 200 μM H_2_O_2_, and the cell viability was measured by MTT assay (Fig. 1). We found that there was no toxicity of H_2_O_2_ up to 50 μM, and the cell growth was inhibited at 100 μM of H_2_O_2_ and completely suppressed at 200 μM. From this result, it was suggested that even the 1.1B4 cells express antioxidant enzymes to survive from H_2_O_2_ stress but at higher concentration above 50 μM of H_2_O_2_, cells were not able to grow well because of not enough antioxidant defense system. Therefore, to find antioxidant system of these cells, comprehensive analysis of gene expression of H_2_O_2_ treated 1.1B4 cells was analyzed by DNA microarray at 100 μM H_2_O_2_. The 1.1B4 cells were treated with 100 μM H2O2 for 4 hours, then total RNA was extracted and labeled with Cy3 then analyzed by Agilent human whole genome microarray. Results were submitted to NCBI GEO data base with accession number GSE83369. (http://www.ncbi.nlm.nih.gov/geo/query/acc.cgi?acc=GSE83369). Total 2903 genes were up-regulated more than two times of untreated cells, and 2283 genes were down regulated less than 1/2. Most of the gene were classified unknown or others, the genes classified to nucleotide metabolism were major group (Table 1, Supplemental Fig. S1, S2). Some of gene expression was confirmed by RT- real time PCR (Table 1). Antioxidant enzyme *HMOX1*, was up-regulated, but *SOD1* and *CAT* were not changed. On the other hand, *GPX1* and *GCLC*, a glutathione synthesis and antioxidant enzymes, were increased. Among the transcription factors that are up-regulated by H_2_O_2_ described by Marinho *et al.* (Marinho et al. 2014), *FOSB* and *JUNB* were strongly up-regulated, suggesting AP-1 pathway was up-regulated. Several apoptosis related enzymes were also up-regulated. Among the antioxidant pathway (Marinho et al. 2014), *KEAP1* was not changed. There were many genes that function was unknown or slightly known, up-regulated or down-regulated. Among them, *NR4A3*, kind of orphan receptor, was up-regulated significantly, therefore, the effect of H_2_O_2_ concentration on the expression of this gene was measured (Fig. 2A). The expression of *NR4A3* was increased by H_2_O_2_ concentration until 100 μM at 4 hours significantly. Therefore, the expression of *NR4A3* after H_2_O_2_ treatment was measured up to 24 hours (Fig. 2B). It is interesting that *NR4A3* expression was increased rapidly and decreased to original level by 24 hours, suggesting that the *NR4A3* is early responsible gene to H_2_O_2_ stress, and it is suggested that NR4A3 might have important role in antioxidant defense system in 1.1B4 cell against H_2_O_2_ stress.

**Fig.1.**
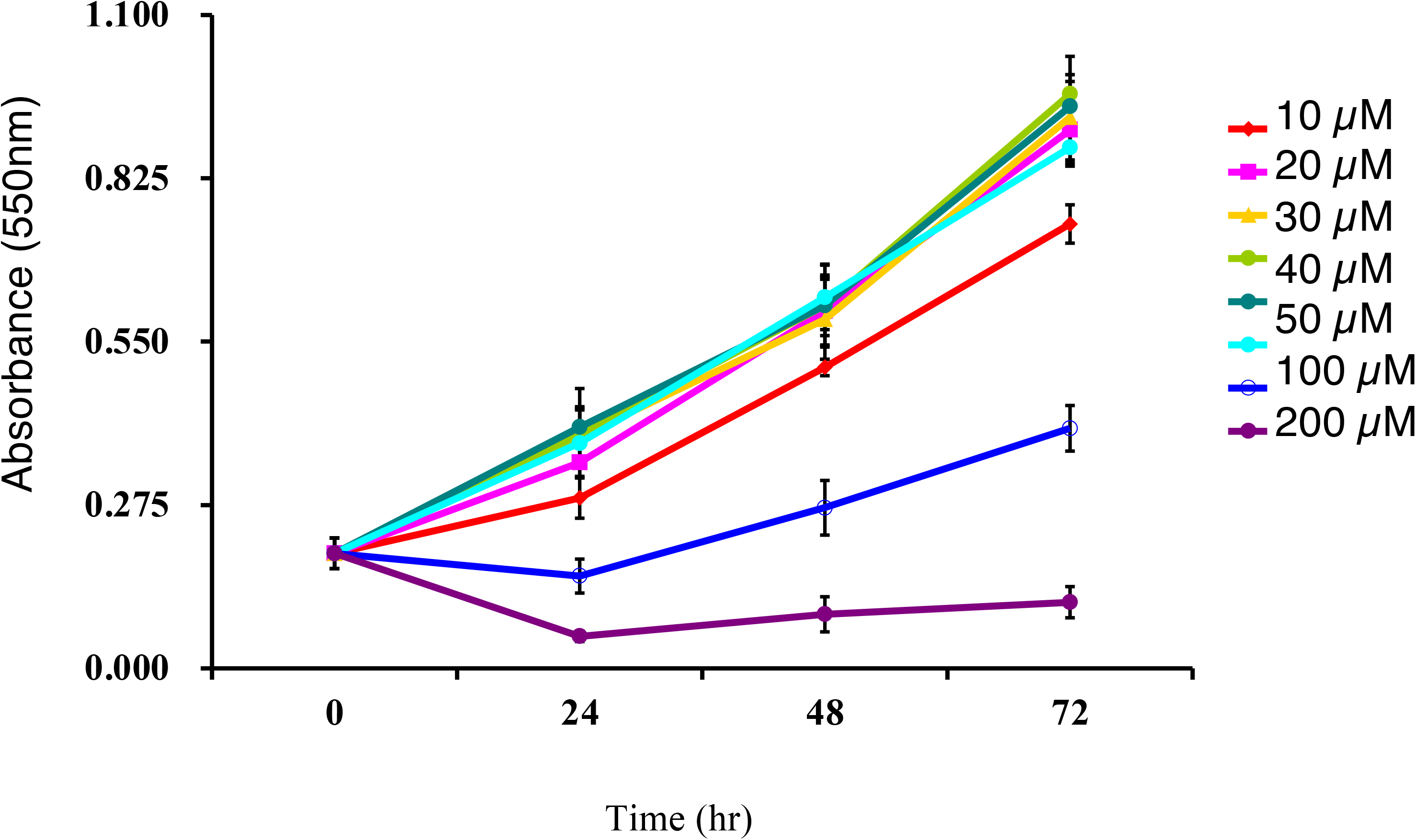
Effect of H_2_O_2_ concentration on the cell growth of 1.1B4 islet derived cell. Islet derived 1.1B4 cell was seeded and cultured in RPMI-1640 medium, and 24 hours later cell was added H_2_O_2_ to 10, 20, 30, 40, 50, 100, 200 μM and cell viability was analyzed at 24, 48 and 72 hours by MTT assay. (n=5)

**Table 1.**
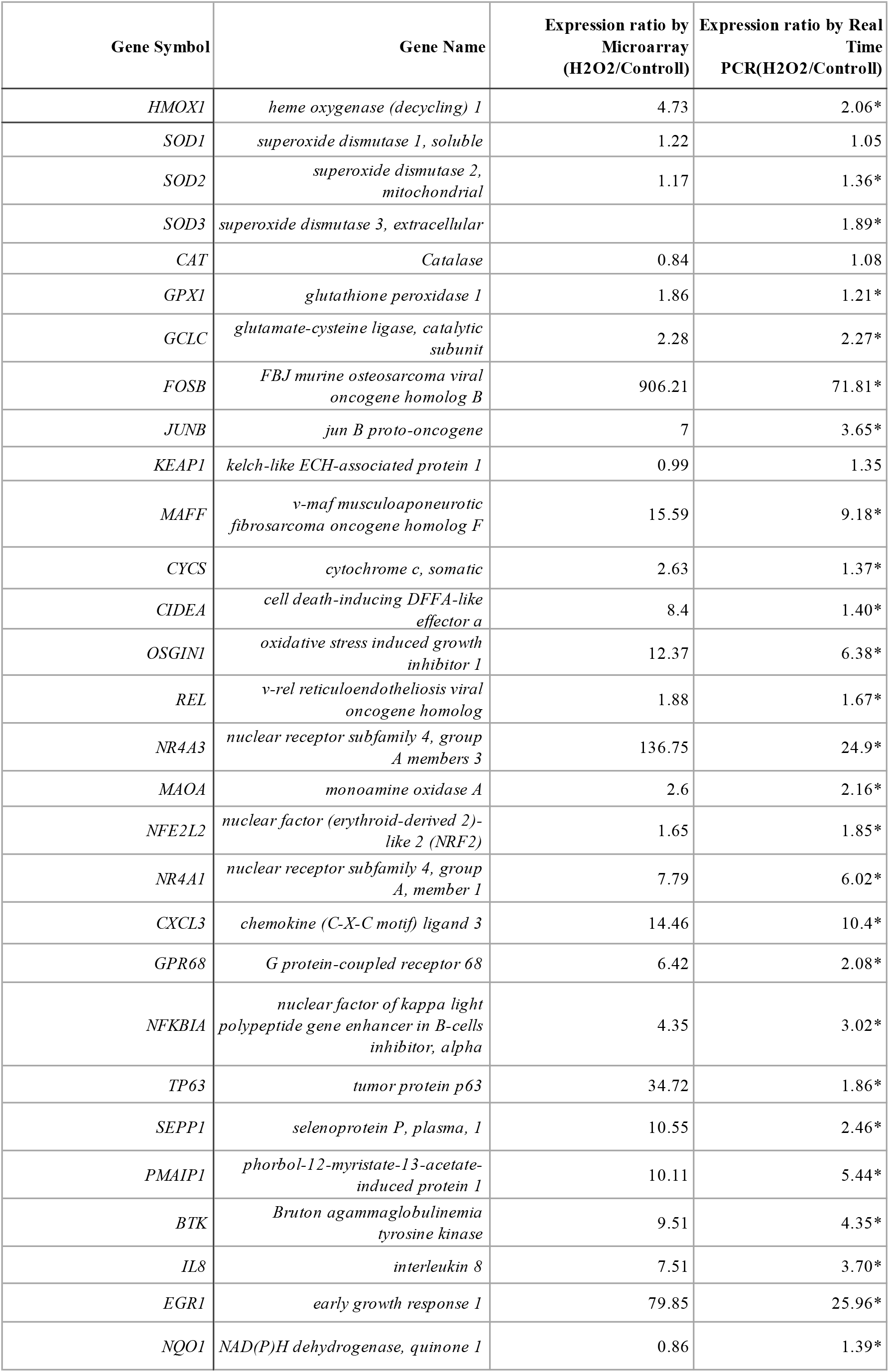
Comparison of expression of genes measured by microarray and real time PCR between H_2_O_2_ treated cells and control cells. Average of 2 microarray data of H2O2 treated cells was compared with 1 microarray data of control cells after normalization. For comparison with real time PCR, 12 RNA from 12 cell culture dishes in 2 groups obtained for microarray were used, and the result of real time PCR was expressed by the average of GAPDH and ß-actin. * p<0.05

**Fig. 2.**
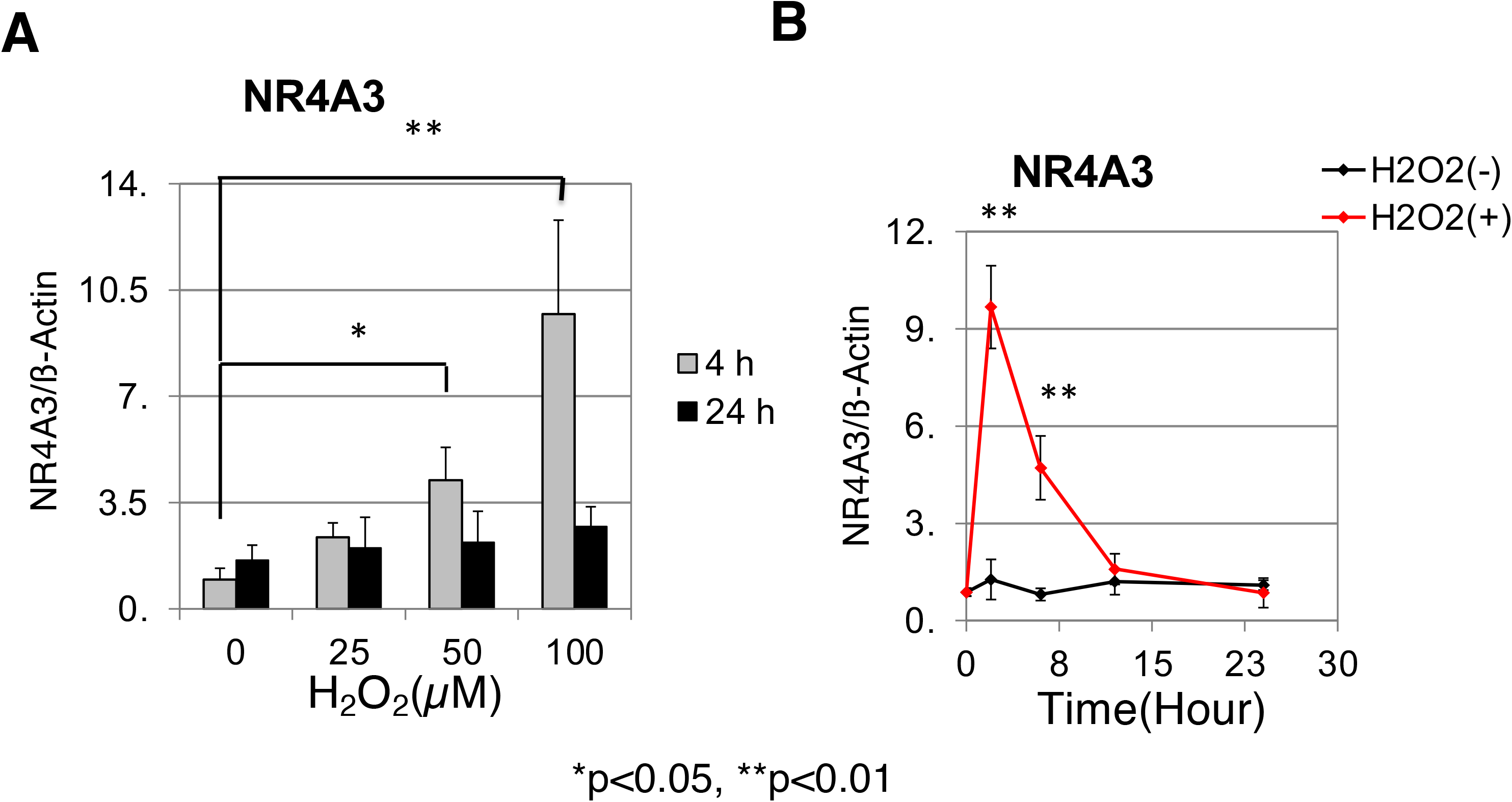
Effect of H_2_O_2_ concentration on *NR4A3* gene expression and time course of expression. **A**, Human 1.1B4 cell (1.0 x 10^5^ cells/ml) was incubated in 2ml culture medium and 24 hours later medium was changed to individual concentration of H2O2 containing medium, then 4 or 24 hours later total RNA was extracted with Isogen. Expression of *NR4A3* was measured by real time PCR with condition B. n=3. **B**, After addition of 100 μM of H_2_O_2_, cell was incubated individual time and RNA was extracted. n=3. ß-Actin was used for control.

### Knock down of *NR4A3* mRNA by RNA interference

To investigate the roll of *NR4A3* in 1.1B4 cell, *NR4A3* mRNA was knocked down with siRNA. The 1.1B4 cells were incubated with 5 nM siRNA of *NR4A3* for 48 hours, the mRNA of *NR4A3* was decreased to 29.6% of control cell (Fig. 3A). The 1.1B4 cells knocked down with siRNA of *NR4A3* were then treated with 100 μM H_2_O_2_, and analyzed cell viability by MTT assay (Fig. 3B). After 72 hours, the viability of siNR4A3 treated 1.1B4 cell was significantly decreased, and when H2O2 was added to culture medium, the growth was completely stopped.

**Fig. 3.**
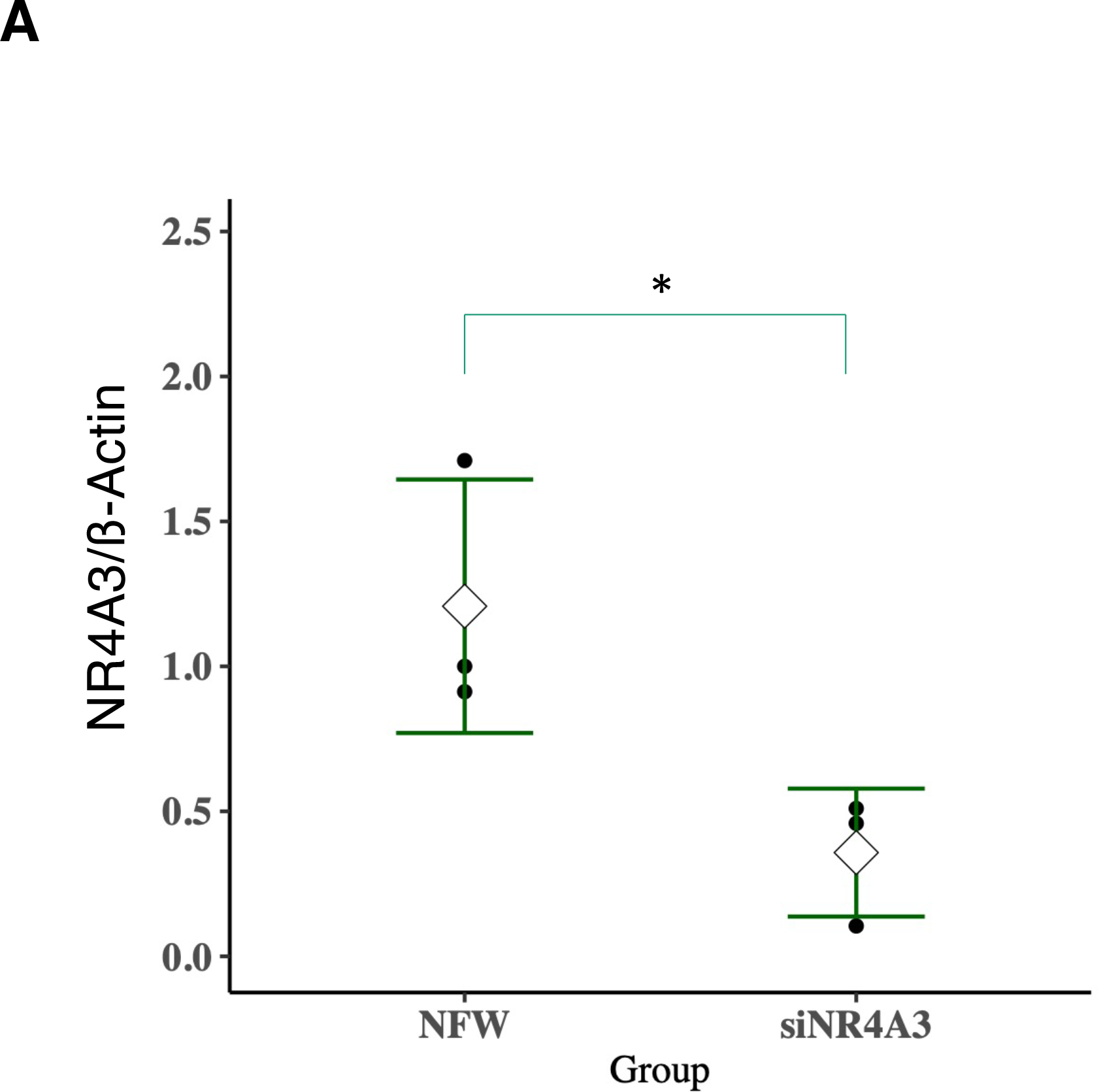
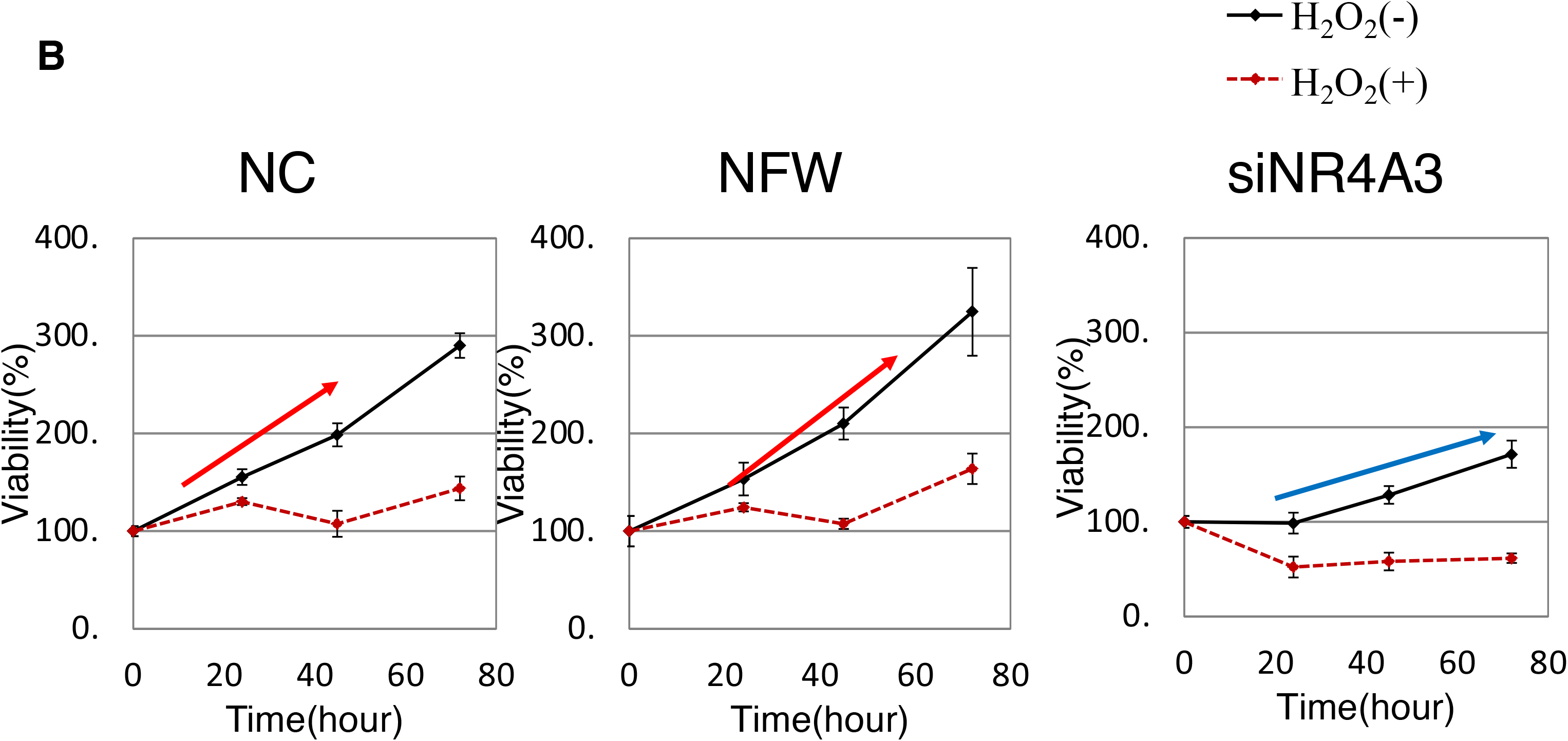
Effect of *NR4A3* knocked down on the expression of *NR4A3* and cell viability. **A**, After knocked down of *NR4A3* gene in 1.1B4 cell with siRNA, total RNA was extracted and expression of *NR4A3* and ß-actin was measured by real time PCR with Cyber green method with condition B. n=3 *p<0.05. **B**, After knocked down of *NR4A3* gene of 1.1B4 cell, cell was incubated with 100 μM of H_2_O_2_, and cell viability was measured by MTT assay. n=5. NC: negative control; NFW: nuclease free water added cell; siNR4A3: siRNA of *NR4A3* treated cell.

### Comprehensive analysis of gene expression in 1.1B4 cells by DNA microarray after the knock down of *NR4A3* with siRNA

To analyze the role of *NR4A3* in pancreatic islet cell further, we performed comprehensive analysis of gene expression of 1.1B4 cells by Agilent human DNA microarray after the knocked down of *NR4A3* mRNA. Data were submitted to NCBI GEO data base with accession number GSE86924. (http://www.ncbi.nlm.nih.gov/geo/query/acc.cgi?acc=GSE86924). We found 1,044 genes were significantly increased over 1.5 folds and 859 genes were significantly decreased under 0.67 folds (Supplemental Fig. S3A, B). Using those genes, pathway analysis was performed by KEGG pathway database. (Supplemental Table S2). Many genes in PI3K-Akt signaling pathway were up-regulated, and Rap1 signaling pathway genes were down-regulated. Changes in gene expression measured by microarray of transcription factor, related to H2O2 addition (Marinho et al. 2014), antioxidant genes and major cell growth related genes are also listed in Table 2. Some of these results were confirmed by real time PCR (Fig. 4A - N, Supplemental Fig. S4A - H). *NR4A3* expression was down regulated (Fig. 4A) but this group of orphan receptor, *NR4A1* (Fig. 4B) was not changed and *NR4A2* (Fig. 4C) was up-regulated. Among the genes of antioxidant enzymes, the expression of *HMOX1* was decreased when measured by microarray, and confirmed by real time PCR (Fig. 4E), the expression of *HMOX1* was reduced to 68%. However, *GCLC* (Supplemental Fig. S4E) was only 0.88 times of control, and *SOD1* (Supplemental Fig. S4F) was not changed significantly. We found interesting gene expression of antioxidant enzyme of *GLRX* (Fig. 4F). After the knocked down the *NR4A3*, *GLRX* expression was increased, different from *HMOX1*. *SOD3* was also increased by *NR4A3* knock down (Fig.4G). Among the transcription factors related to H2O2 oxidation, *MAFA* was slightly down regulated measured by microarray but was not changed measured by real time PCR (Supplemental Fig. S4A - D). Other redox sensitive transcription factors were not down regulated.

**Table 2.** Changes of gene expression after siRNA of NR4A3 treatment measured by microarray. Average of 3 microarray data of siRNA of NR4A3 treated cells after normalization was compared with average of 3 microarray data of control cells.

| Gene symbol | Gene name | Fold changes<br>(SiRNA vs<br>Cont) | Gene<br>symbol | Gene name | Fold changes<br>(SiRNA vs<br>Cont) |
| --- | --- | --- | --- | --- | --- |
| **Orphan receptor** |  |  | **Cell cycle** |  |  |
| NR4A3 | nuclear receptor subfamily 4, group A members 3 | 0.85 | E2F1 | E2F transcription factor 2 | 0.66688 |
| NR4A2 | nuclear receptor subfamily 4, group A, member 2 | 1.06 | CDK1 | cyclin-dependent kinase 1 | 0.7342 |
|  |  |  | CDK2 | cyclin-dependent kinase 2 | 0.34304 |
| **Antioxidant responsible Transcription Factor** |  |  | CDK4 | cyclin-dependent kinase 4 | 0.87803 |
| MAFA | v-maf musculoaponeurotic fibrosarcoma oncogene homolog A (avian) | 0.75 | CCNA1 | cyclin A1 | 0.84123 |
| FOSB | FBJ murine osteosarcoma viral oncogene homolog B | 0.96 | CCNB1 | cyclin B1 | 0.72389 |
| FOS | FBJ murine osteosarcoma viral oncogene homolog | 2.5 | CCND1 | cyclin D1 | 0.99547 |
| JUNB | jun B proto-oncogene | 1.02 | CCNE1 | cyclin E1 | 0.44222 |
| JUN | jun proto-oncogene | 0.76 | CCNY | cyclin Y | 0.62658 |
| KEAP1 | kelch-like ECH-associated protein 1 | 0.93 | CDKN1B | cyclin-dependent kinase inhibitor 1B (p27, Kip1) | 1.66505 |
| NFE2L2 | nuclear factor (erythroid-derived 2)-like 2, NRF2 | No data | CDKN1B | cyclin-dependent kinase inhibitor 1B (p27, Kip1) | 1.66505 |
| CREB5 | cAMP responsive element binding protein 5 | 2 |  |  |  |
| TP53 | tumor protein p53 | 0.83 | **Others** |  |  |
| NOTCH4 | notch 4 | 1.36 | RAB5B | member RAS oncogene family (RAB5B) | 0.54527 |
| NFKB1 | nuclear factor of kappa light polypeptide gene enhancer in B-cells 1 | 0.95 | RAP1B | RAP1B, member of RAS oncogene family | 0.58507 |
| NFKB2 | nuclear factor of kappa light polypeptide gene enhancer in B-cells 2 (p49/p100) | 0.77 | TGFB3 | transforming growth factor, beta 3 | 0.50684 |
| SP1 | Sp1 transcription factor | 0.8 | PIK3CA | phosphoinositide-3-kinase, catalytic, alpha polypeptide | 1.24095 |
| HIF1A | hypoxia inducible factor 1, alpha subunit (basic helix-loop-helix transcription factor) | 1.02 | MRAS | muscle RAS oncogene homolog | 0.4872 |
| SREBF1 | sterol regulatory element binding transcription factor 1 | 0.76 | RAP1B | RAP1B, member of RAS oncogene family | 0.59426 |
| HSF1 | heat shock transcription factor 1 | 0.84 | SREBF1 | sterol regulatory element binding transcription factor 1 (A_33_P3222139) | 0.85885 |
|  |  |  | HIF1A | hypoxia inducible factor 1, alpha subunit (basic helix-loop-helix transcription factor) | 0.91634 |
| **Antioxidant Responsible Enzyme** |  |  | BAD | BCL2-associated agonist of cell death | 0.65799 |
| HMOX1 | heme oxygenase (decycling) 1 | 0.48585 | AKT1 | v-akt murine thymoma viral oncogene homolog 1 | 0.85096 |
| GCLC | glutamate-cysteine ligase, catalytic subunit | 0.88148 | BAD | BCL2-associated agonist of cell death | 2.09416 |
| GPX1 | glutathione peroxidase 1(A_33_P3239849) | 0.81867 | FOXO1 | Homo sapiens forkhead box O1 (FOXO1), mRNA [NM_002015] | 1.0298 |
| GPX1 | glutathione peroxidase 1(A_33_P3354322) | 0.81248 | FOXO3 | Homo sapiens forkhead box O3 (FOXO3), transcript variant 1, mRNA | 1.08933 |
| CAT | catalase | 1.59628 | GAPDH | glyceraldehyde-3-phosphate dehydrogenase | 0.73695 |
| SOD1 | superoxide dismutase 1, soluble | 0.87111 | TBP | TATA box binding protein | 0.93576 |
| SOD2 | superoxide dismutase 2, mitochondrial | 0.6454 | ACTB | actin, beta | 0.89854 |
| SOD3 | superoxide dismutase 3, extracellular | 0.65966 |  |  |  |
| GLRX | glutaredoxin (thioltransferase) | 1.97201 |  |  |  |
| GLRX2 | glutaredoxin 2 | 0.83005 |  |  |  |
| TXN | thioredoxin | 0.82374 |  |  |  |
| PMEL | premelanosome protein | 1.6221 |  |  |  |

**Fig. 4.**
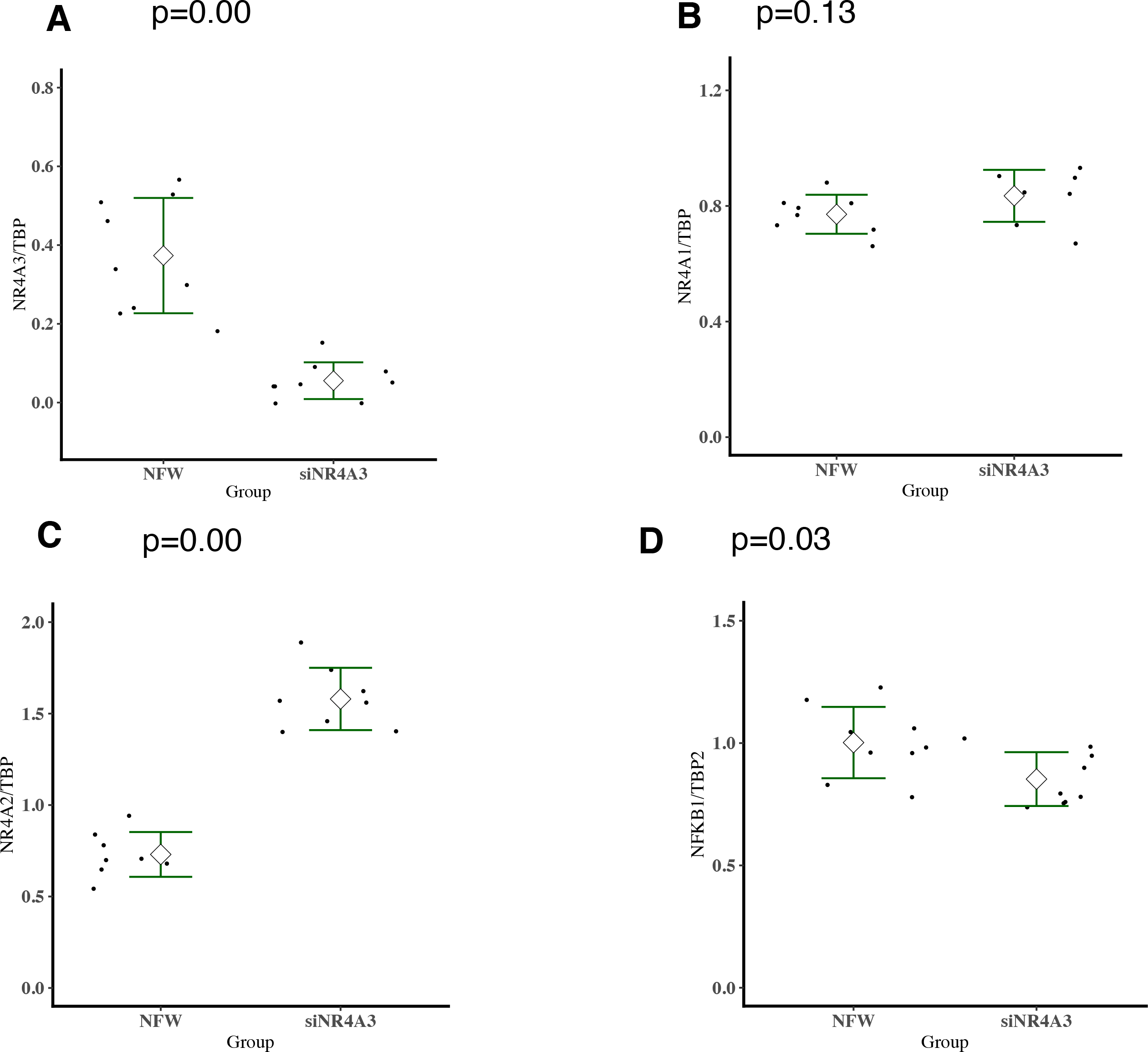
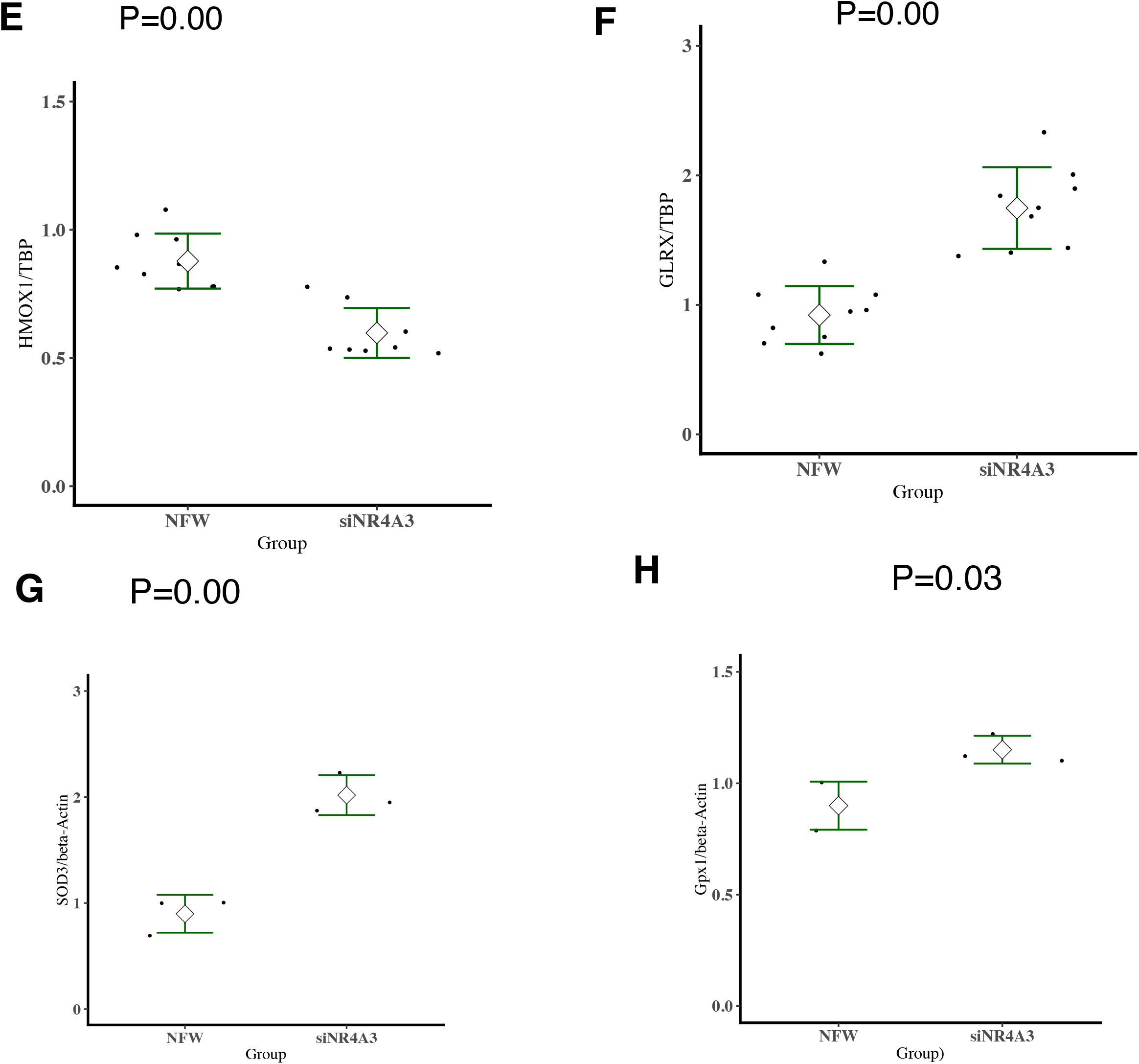
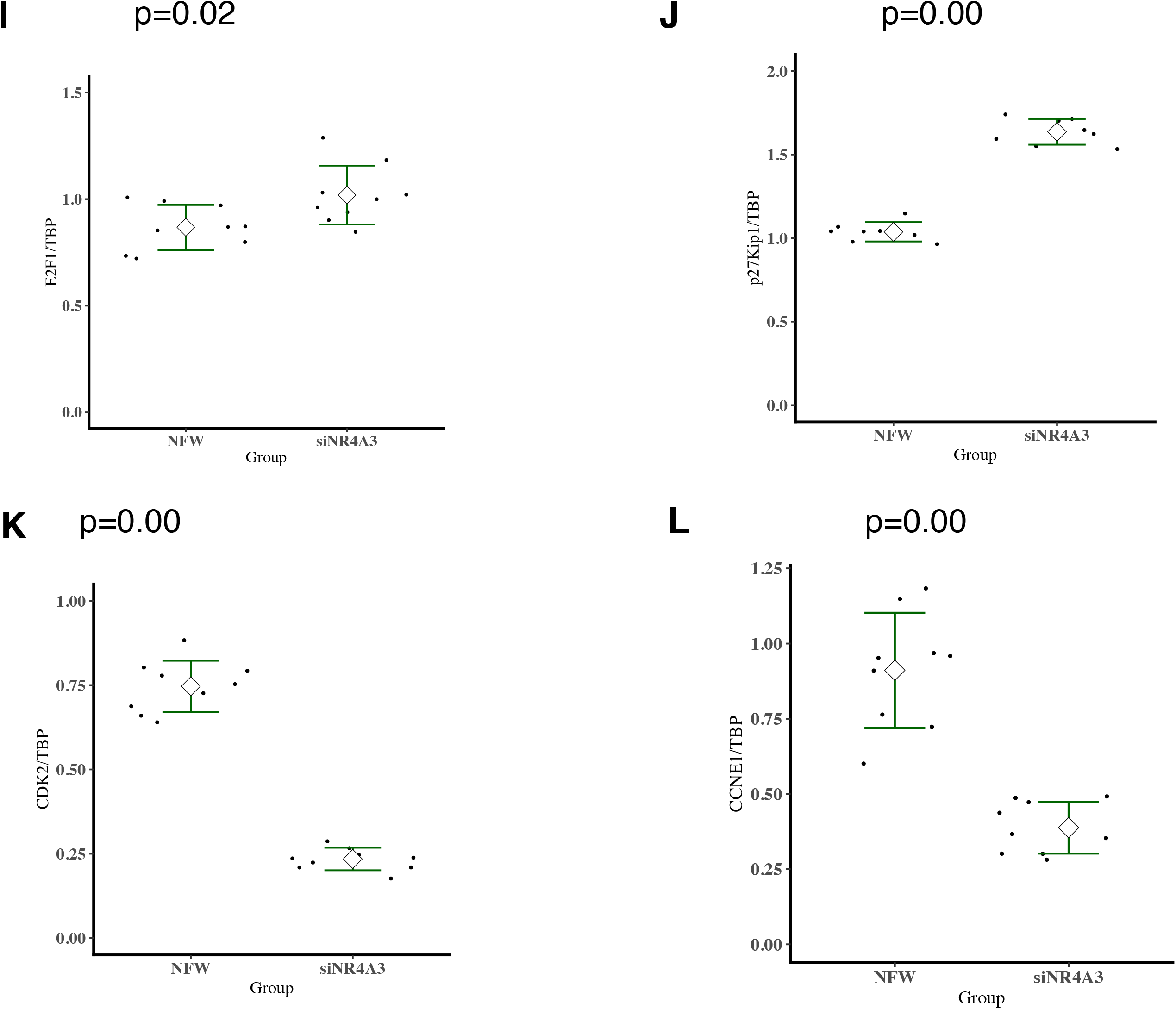
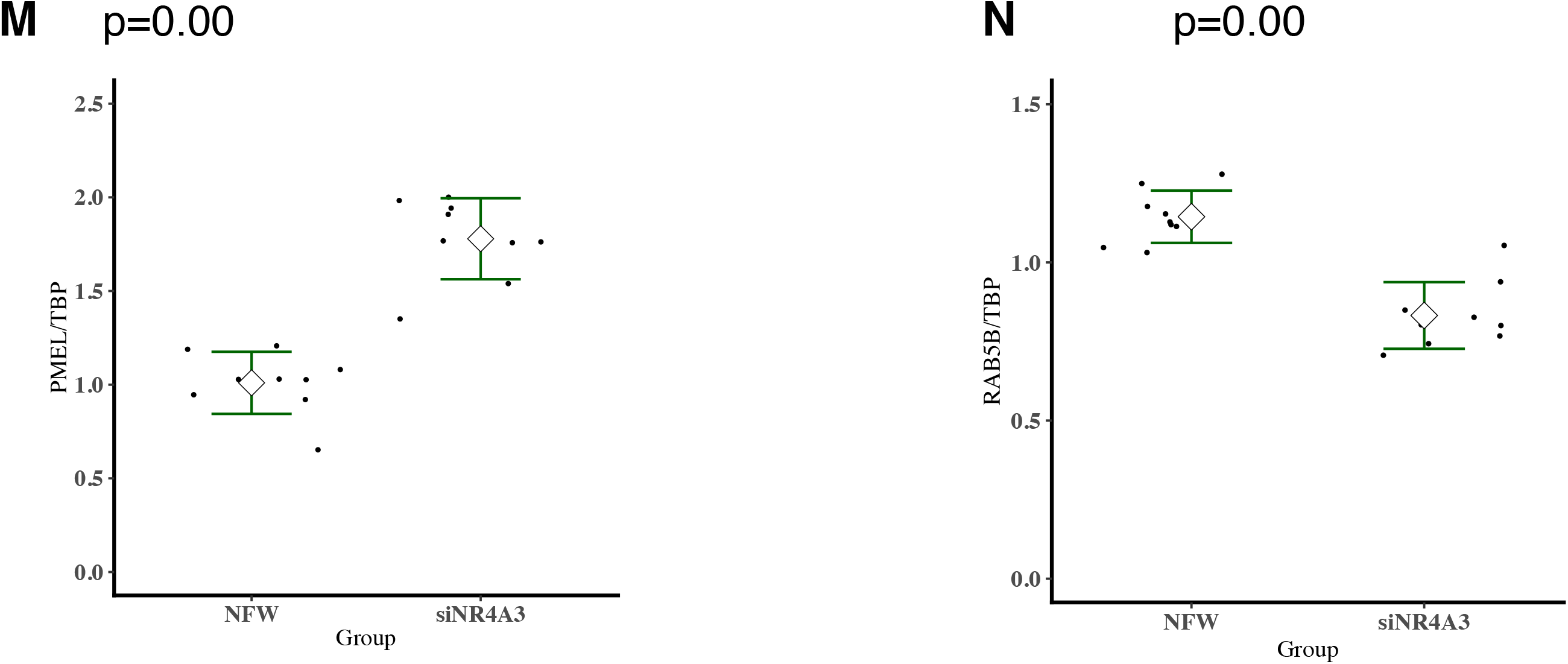
Changes of Various Gene expression in 1.1B4 Cell after the knocked down of *NR4A3* mRNA measured by real time PCR. (A) Human 1.1B4 cells were incubated 24hours, half of them (n=9) were added siRNA (siNR4A3) and another half (n=9) were added nuclease free water (NFW), and after 52 hours, mRNA was purified with RNeasy Plus Mini Kit. Expression of *NR4A3* was measured by real time PCR with Taqman method using condition C. TATA binding protein (TBP) mRNA was used for internal control. (B) The *NR4A1* expression, (C) *NR4A2* expression. (D) - (E) *NFKB1*and *HMOX1* respectively. (F) *GLRX* expression (n=8). (G) Human 1.1B4 cells were incubated 24hours, half of them (n=3) were added siRNA (siNR4A3) and another half (n=3) were added NFW, and after 52 hours, mRNA was purified with Isogen reagent and treated with RNase-free DNase. Expression of *SOD3* was measured by real time PCR with Cyber green method using condition B. β-Actin was used for control. (H) Expression of *GPX1* using same method of (G). (I) *E2F1* using same method of (A) (n=8). (J) - (M) *p27Kip1*, *CDK2*, *CCNE1* and *PMEL*, respectively, using same method of (A). (N) *RAB5B* using same method of (A) (n=8).

Gizard *et al.* (Gizard et al. 2011) reported that NOR1 (NR4A3 or Nur77) is recruited to a nerve growth factor-induced clone B response element (NBRE, aaaggaca or aaaggtca) site (Philips et al. 1997). Therefore, whether the *HOMX1* gene has NBRE sequence, or not, was investigated. The *HMOX1* gene structure was obtained from NCBI database (AY460337). There are two NBRE like sequence of aaggtca, tentative NR4A3 responsible element, at second intron (Fig. 5). Transcription factor responsible element in second intron might works when H2O2 is present in 1.1B4 cell culture medium.

**Fig. 5.**
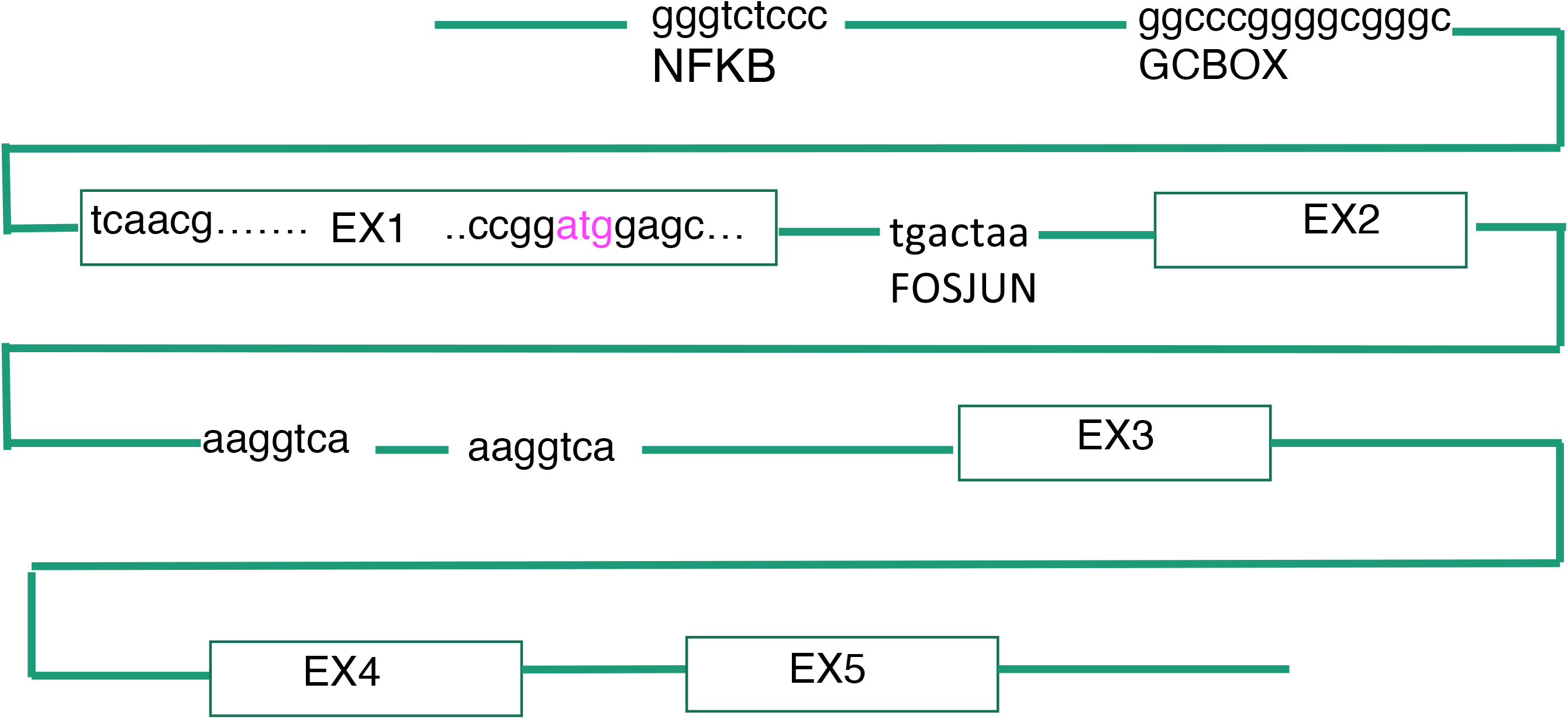
Gene structure of *HMOX1*. Human *HMOX1* gene was obtained from PubMed gene (ACCESSION AY460337). NR4A3 binding motif of aaggtca were searched and abstract of the gene structure was displayed. NFkB shows NFkB binding motif. FOSJUN shows FOS and JUN binding motif. EX1 *etc*. shows exson 1 *etc*.

As described before, knock down cells decreased cells growth, therefore, cell cycle regulation genes that expression was modified by siRNA of *NR4A3*, were investigated. *CCNE1* (*Cyclin E*) and *CDK2* expressions were decreased remarkably (Table 2). Then those down-regulations were confirmed by RT-real time PCR (Fig. 4K, L). *CDK2* and *CCNE1* expression were decreased strongly compared with un-treated cell. To investigate the mechanism, gene structure of *CDK2* and *CCNE1* were analyzed (Supplemental Fig. S5A, B and S6). There was no aaggtca at up-stream of *CDK*2, however, down-stream of *CDK2*, there is aaggtca sequence. This area is also up-stream of the gene of *RAB5B*, therefore, the effect of siRNA of *NR4A3* on this *RAB5B* expression was investigated. There is also *PMEL* is near this area, the expression of *PMEL* was also measured. As shown in Fig 4M and N, those two genes expression were also decreased or increased by siRNA of *NR4A3*. It is, therefore, suggested that *NR4A3* promotes proliferation through activating *Rap1* signaling pathway, therefore, it is also suggested that the *NR4A3* contribute to homeostasis against extracellular stress such as oxidative stress.

## DISCUSSION

It is known that pancreatic islet ß-cell weakly express antioxidant enzymes, therefore, oxidative stress enhances diabetes mellitus. As shown in this experiment of microarray of H_2_O_2_ treated 1.1B4 cell, among the antioxidant enzymes, *HMOX1, GPX1* and *GCLC* were up-regulated (Table 1). Other major antioxidant enzymes were not up-regulated, for example, *SOD1* and *CAT* were not induced by H_2_O_2_ addition, therefore, those *HMOX1, GPX1* and *GCLC* enzymes are responsible for antioxidant defense system in pancreatic cell, and those are very important enzymes for defending against oxidative stress at diabetes. Alam and Cook (Alam and Cook 2007) described that there are 4 pathways that regulates *HMOX1* gene expression at oxidative stress. But here, not only those 4 pathway, we showed that *HMOX1* and *NR4A3* were up-regulated when H_2_O_2_ was added, and the expression of *HMOX1* was down-regulated when *NR4A3* was knocked down by siRNA resulting loss of antioxidative resisstance. Furthermore, the second intron of *HMOX1* has NBRE like sequence of aaggtca. Therefore, we concluded that the *NR4A3* is antioxidant responsible transcription factor and regulates *HMOX1* expression at oxidative stress. Not only 1.1B4 cell, *NR4A3* expression was also increased in HUCF2 cell at oxidative stress (Shimizu et al. 2015). Therefore, we propose that the *NR4A3* is new oxidative stress responsible transcription factor not listed before (Marinho et al. 2014), and this pathway is major antioxidant pathway in 1.1B4 cell (Graphical abstract). *FOSB* is also up-regulated by H_2_O_2_ in this experiment, therefore, FOS pathway might be also another major antioxidant pathway.

There are two NBRE like sequence of aaggtca in *HMOX1*. NBRE contains aggtca which is typically recognized by retinoic acid receptor/retinoid X receptor (RAR/RXR) subfamily, and it also includes two aa residues preceding this hexanucleotide (Philips et al. 1997). On the other hand, NR4A1 and NR4A2 forms heterodimer with RXR and binds to direct repeat of aggtca (Safe et al. 2016). Therefore, we propose aaggtca is NR4A3 responsible element.

Micro-array analysis has shown that knock down of *NR4A3* decreased *CDK2* and *CCNE1* (*cyclin E*) expression (Table 2). This result confirmed by real time PCR and promotor region analysis (Fig. 4K and 4L, Supplemental Fig. S5A, B and S6). These results show that *NR4A3* is key transcription factor not only for antioxidant system but also for cell cycle control in 1.1B4 cell. Tessem *et al.* (Tessem et al. 2014) demonstrate that *E2F1* and *CCNE1* are key cell cycle inducers. Our result of *cyclin E* consistent with Tessem results, but our results showed *E2F1* expression was only slightly increased in *NR4A3* knock down cell (Fig. 4I). Therefore, it is suggested that NR4A3 directly controls *cyclin E* and *CDK2,* not via *E2F1*.

Robertson (Robertson 2010) described that main stays of therapy for type 2 diabetes involve drugs that are insulin-centric, i.e., they are designed to increase insulin secretion and decrease insulin resistance. The mechanism for this unrelenting deterioration of ß-cell function is related to chronic oxidative stress. This suggests that drug discovery should not exclusively focus on insulin-centric targets, but also include glucose-centric strategies, such as antioxidant protection of the ß-cells. This may facilitate repair of ß-cells undergoing damage by oxidative stress secondary to chronic hyperglycemia. Furthermore, Gao *et al*. (Gao el al. 2014) have shown that over expression of *NR4A3* results in down-regulation of insulin gene transcription and insulin secretion. It is suggested that oxidative stress up-regulated *NR4A3* expression resulting down-regulation of insulin secretion. Therefore, antioxidant material might recover from islet damage. For this aim, many antioxidants were used for diabetes therapy. For example, Cinnamtannin D-1, one of the main A-type procyanidin oligomers in *C. tamala*, was discovered to dose-dependently reduce palmitic acid- or H_2_O_2_-induced apoptosis and oxidative stress in INS-1 cells, MIN6 cells, and primary cultured murine islets (Wang et al. 2014), however, antioxidant drugs were not in use for human diabetes. Therefore, we screened new antioxidant from food stuff for potential therapy use, and recently found strong antioxidant Zeylaniin A from edible vegetable (Nomi et al. 2012). This polyphenol may be used for diabetes treatment.

For microarray experiment, Agilent array was used and confirmed by real time PCR. Some times other type of microarray shows different data from real time PCR and shows false positive data, but Agilent array used here shows very co-related data to real time PCR as far as we measured in this experiment. And three microarray data of same conditions showed very similar data. Use of internal control gene for real time PCR was not easy, because many house keeping genes are known to change their transcription depend on the conditions. Therefore, average of two house keeping genes of *GAPDH* and *β*-actin were used for H_2_O_2_ experiment. *TBP* was also used for siRNA treated cell because *TBP* was not changed by siNR4A3 treatment measured by microarray (Table 2).

## METHODS

### Reagents

RPMI-1640 medium (GIBCO, Tokyo or Sigma, Tokyo), FBS (CELLect FBS, MP Bomedical Japan, Tokyo Japan, LOT#7997K or Biowest, Nuaillé, France, Lot.No:S05831S1820), Penicilin (50 IU/ml) - Streptomycine (50 micro g/ml) ( ICN Biomedicals, Irvine, California, United States), 96 well plate (IWAKI, Tokyo, Japan), NR4A3 Silencer Select siRNA (Applied Biosystems - Thermo Fisher Scientific Japan, Tokyo, siRNA ID: s15542), Silencer Select Negative Control #1 siRNA, Nuclease Free Water and Lipofectamine 2000 (Applied Biosystems - Thermo Fisher Scientific Japan, Tokyo). RNeasy Plus Mini Kit (Qiagen, Tokyo, Japan), Bio-analyzer (Agilent Technology, Palo Alto, CA, USA). Isogen reagent and RNase-free DNase (Nippon gene, Toyama, Japan). Agilent Whole Human Genome Oligo Microarray 4×44K Ver. 2.0 (Agilent Technology, Palo Alto, CA, USA) Reagents for microarray (Agilent Technology, Palo Alto, CA, USA). Agilent G2565BA Microarray Scanner System (Agilent Technology, Palo Alto, CA, USA). The scanned images were analyzed with Feature Extraction Software 9.5.1.1 (Agilent Technology, Palo Alto, CA, USA) using default parameters (protocol GE1-v5_95_feb07 and Grid: 014850_D_F_20101031). Spot Fire software (TIBCO, NTTCom, Tokyo, Japan) and the GeneSpringGX10 v 7.3.1 (Agilent Technology, Palo Alto, CA, USA). DNaseI-treated total RNA with M-MLV reverse transcriptase (Invitrogen, Carlsbad, CA, USA) and Oligo (dT) 15 primer (Promega, Madison, WI, USA). STEP ONE PLUS Real Time PCR system (Applied Biosystems Japan, Tokyo, Japan), SYBR GREEN PCR Master Mix (Applied Biosystems Japan, Tokyo, Japan). Takara Premix Ex Taq (Probe qPCR) (TaKaRa, Kyoto, Japan), Designs of PCR probes: Universal Probe Library Assay Design Center at Roche (http://www.universalprobelibrary.com). ABI PRISM 7900HT Sequence Detection System (Applied Biosystems, Tokyo, Japan).

### Biological Resources: Human pancreatic islet derived cell, 1.1B4

Islet derived hybrid cell of 1.1B4 (ECACC No. 10012801) formed by the electrofusion of a primary culture of human pancreatic islets with PANC-1, a human pancreatic ductal carcinoma cell line (ECACC catalogue number 87092802), was obtained from DS Pharma Biomedical Co., Ltd., Osaka, Japan, and cultured in RPMI-1640 medium (GIBCO, Tokyo or Sigma, Tokyo, Japan) supplemented with 10% FBS (CELLect FBS, MP Bomedical Japan, Tokyo Japan, LOT#7997K or Biowest, Nuaillé, France, Lot.No:S05831S1820) with Penicilin (50 IU/ml) - Streptomycine (50 μg/ml) ( ICN Biomedicals, Irvine, CA, USA) in humidified air at 37 °C with 5% CO_2_.

### Cell toxicity of H_2_O_2_ and the effect of H_2_O_2_ on gene expression of 1.1B4

The cells of 1.1B4 were cultured in 96 well plate (IWAKI, Tokyo, Japan) at 6.0 x 10^3^ cells / well with RPMI-1640 medium for 24 hours, then 0, 10, 20, 30, 40, 50, 100, 200 μM H_2_O_2_ was added to medium, and cell viability was analyzed at 24, 48 and 72 hours by MTT (3-(4,5-dimethylthiazol-2-yl)-2,5-diphenyltetrazolium bromide) colorimetric assay (Mosmann 1983). We found that there was no toxicity up to 150 μM H_2_O_2_. Next, we examined the effect of H_2_O_2_ on gene expression by real time PCR. Human 1.1B4 cell (1.0 x 10^5^ cells/ml) was incubated in 2ml culture medium and 24 hours later medium was changed to 25, 50 and 100 μM H_2_O_2_ containing medium, then 4 or 24 hours later total RNA was extracted with Isogen. Expression was measured by real time PCR with condition B. Cells of 1.1B4 were also treated with 100 μM H_2_O_2_ and we analyzed gene expression after the exposure to H_2_O_2_ for 2, 6, 12, 24 hours.

### Knock down of NR4A3 mRNA by RNA interference

NR4A3 Silencer Select siRNA (ABI, siRNA ID: s15542) was used for knocked down the *NR4A3* mRNA, and Silencer Select Negative Control #1 siRNA (ABI) and Nuclease Free Water (ABI) were used for negative control. *NR4A3* mRNA of 1.1B4 cells was knocked down by incubating with 5 nM siRNA and 0.1 % of Lipofectamine 2000 for 48 h. Total RNA was isolated by Isogen then mRNA was measured as described before. The 1.1B4-siNR4A3 treated cells or control cells were treated with or without 100 μM H_2_O_2_, and cell viability was assayed by MTT assay.

### Extraction and purification of mRNA

For micro-array, total RNA was isolated using the RNeasy Plus Mini Kit (Qiagen, Tokyo, Japan). The purity of RNA was assessed by a Bio-analyzer (Agilent Technology, Palo Alto, CA, USA) before microarray analysis. The RNA thus obtained was also used for real time PCR. For only real time PCR use, total RNA was extracted with Isogen reagent (Nippon gene, Toyama Japan) as described previous paper (Oba et al. 2006) and treated with RNase-free DNase (Nippon gene, Toyama, Japan).

### Global analysis of gene expression by DNA microarray in 1.1B4 cells

(**Experiment A**) Pancreatic derived 1.1B4 cell was seeded as 1.5×10^5^ cells / ml to RPMI-1640 medium (GIBCO, Tokyo, Japan) in 6 well plate, and 24 hours later, cell was incubated with 100 μM H_2_O_2_ 4 hours, then total RNA was extracted with RNeasy Mini Kit (Qiagen, Tokyo) (n=6 for control and treated respectively). The quality of all 12 RNA samples was checked by Agilent RNA 6000 Nano Reagents Part1 (Agilent Technologies, Tokyo, Japan), and 3 samples of RNA were combined to one microarray sample, then analyzed by Agilent human microarray by the protocol of manufacture. In brief, Cy3-labeled cRNA was generated from 200 ng input total RNA using Agilent’s Low Input Quick Amp Labeling Kit (Agilent Technologies, Tokyo, Japan). cDNA was generated with the primer containing a T7 polymerase promoter, and then was transcripted into cRNA in accompany with the dye labeling. For every type of cell, 1.65 μg cRNA from each labeling reaction was hybridized to the Agilent Whole Human Genome Oligo Microarray (Agilent Technologies, Tokyo, Japan). The Whole Human Genome Oligo Microarray is in a 4 x 44k slide format and each block represents more than 41,000 unique human genes and transcripts. After hybridization, the slides were washed and then scanned with the Agilent G2565BA Microarray Scanner System (Agilent Technologies, Tokyo, Japan). The scanned images were analyzed with Feature Extraction Software 9.5.1.1 (Agilent Technologies, Tokyo, Japan) using default parameters (protocol GE1-v5_95_feb07 and Grid: 014850_D_F_20101031) to obtain background subtracted and spatially detrended Processed Signal intensities. Features flagged in Feature Extraction as Feature Non-uniform outliers were excluded. Data were further normalized using Gene Spring GX10 using default parameters of recommended protocol with Median shift normalization to the 75th percentile and baseline transformed to the median of all samples. Data normalization and filtering was performed using Spot Fire software (TIBCO, NTTCom, Tokyo, Japan) and the Gene Spring v 7.3.1 (Agilent Technologies, Tokyo, Japan). After the reading, one of microarray for control sample was scratched, therefore, the gene expression was compared average of two treated array samples vs one control sample. These 12 RNA samples were also used for real time PCR analysis.

(**Experiment B**) The microarray experiment was designed as follows. Human 1.1B4 cells (n=24) were incubated in RPMI-1640 medium 24hours, half of them (n=12) were added siRNA (siNR4A3) and another half (n=12) were added nuclease free water (NFW), and 48 hours later 3 cells of each group were added H_2_O_2_ to 100 mM. Those 6 cells samples were stored but not used for this paper. After 52 hours, mRNA of remaining 18 cells was purified with RNeasy Plus Mini Kit. The quality of mRNA was checked by Bio-analyzer, 3 mRNA samples were combined to one microarray sample (3 knocked down microarray samples and 3 control microarray samples), and applied on microarray analysis with Agilent Whole Human Genome Oligo Microarray 4 x 44k. The expression was analyzed by Gene-spring software and Spot fire software. Average of 3 array samples treated by H2O2 was compared to average of 3 array samples without treated by H_2_O_2_. For real time PCR, 18 RNA samples were used individually. Real time PCR with Taqman method was used (condition C). *TBP* mRNA was used for internal control. All the sample-labeling, hybridization, washing, and scanning steps were conducted following the manufacturer’s specifications as described before.

### Real-time RT-PCR

The cDNAs were synthesized from DNaseI-treated total RNA with M-MLV reverse transcriptase (Invitrogen, Carlsbad, CA, USA) and Oligo (dT) 15 primer (Promega, Madison, WI, USA) according to the manufacturer’s instructions. Analysis of gene expression was performed with a STEP ONE PLUS Real Time PCR system (Applied Biosystems Japan, Tokyo, Japan) using SYBR GREEN PCR Master Mix (Applied Biosystems Japan, Tokyo, Japan) and the primers listed in Supplemental Table S1. **Condition A**, 0.2 μl of cDNA was mixed with 0.1 μl forward primer (50 pmol / ml), 0.1 μl reverse primer (50 pmol / ml), 4.7 μl water and 5 μl 2×SYBR Green PCR Master MIX. Reactive condition was first incubated at 50 °C for 2 min and 95 °C for 10 min, followed by 40 cycles at 95 °C for 15 sec and 60 °C for 1 min, then final of 95 °C for 15 sec, 60 °C for 15 sec and 95 °C for 15 sec. Normalization of the data was achieved by quantitating the cycle time at an arbitrary fluorescence intensity in the linear exponential phase using Step One Plus Real-Time system Software (Applied Biosystems Japan, Tokyo, Japan) by calculating the ratio of the relative concentration of each mRNA relative to that of average of *GAPDH* and *ß-actin* or *ß-actin* only. To confirm amplification specificity, the PCR products from each primer pair were subjected to a melting curve analysis. The relative quantification of gene expression was computed by using the comparative Ct (threshold cycle) method. **Condition B**, 0.2 μl of cDNA was mixed with 1 μl forward primer (50 μM), 1 μl reverse primer (50 μM), 2.8 μl water and 5 ml 2×SYBR Green PCR Master MIX with same reaction condition.

**Condition C** was performed with a TaqMan method. Briefly, the synthesized cDNA products were subjected to real-time PCR in a reaction mixture (10 μl) containing Takara Premix Ex Taq (Probe qPCR) (TaKaRa, Kyoto, Japan), 200 nM hydrolysis probes and 200 nM PCR primers. Two μl of cDNA was mixed with 5 μl of Premix Ex Taq (2×conc.), 0.2 μl of ROX Reference Dye (50X), 0.2μl of forward primer (10 μM), 0.2 μl of reverse primer (10 μM) and 0.2 μl of TaqMan Probe (10 μM). Designs of PCR probes and primers were obtained from the Universal Probe Library Assay Design Center at Roche (http://www.universalprobelibrary.com) (Supplemental Table S1). Real-time amplifications were performed on the ABI PRISM 7900HT Sequence Detection System (Applied Biosystems, Tokyo, Japan). The settings for the thermal profile were an initial denaturation (30 s at 95°C) followed by 40 amplification cycles: denaturation for 5 s at 95°C; annealing for 30 s at 60°C. Gene-specific standard curves were generated using 2-fold serial dilutions of cDNA. The amount of target mRNA was expressed as the ratio to *TBP* mRNA. Primers are listed in Table S1.

### Classification of genes

The genes up-regulated or down-regulated significantly were categorized by panther process name of the gene prepared by Agilent data for human microarray.

### Pathway analysis and statistical analysis

The genes up-regulated significantly were submitted to the KEGG pathway analysis at Kyoto University. Statistical analysis was performed with R (R Core Team; R: A language and environment for statistical computing. R Foundation for Statistical Computing, Vienna, Austria; 2017 URL https://www.R-project.org/). For comparisons between groups, analysis of covariance (ANNOV method and t test for post hoc test) was used to assess the statistical significance of mean differences between groups, and pairwise t test was used for post hoc testing. The significance levels were set at Pr(>F) 0.05 for anova, and p < 0.05 and p < 0.01 for post hoc testing.

### DATA ACCESS

Microarray data of H2O2 treated cells is available in NCBI GEO data base with accession number GSE83369. (http://www.ncbi.nlm.nih.gov/geo/query/acc.cgi?acc=GSE83369).

Microarray data of siRNA of NR4A3 treated cells is available in NCBI GEO data base with accession number GSE86924. (http://www.ncbi.nlm.nih.gov/geo/query/acc.cgi?acc=GSE86924).

## COMPETING INTEREST STATEMENT

## CONFLICT OF INTEREST

No potential conflicts of interest relevant to this article were reported. Declarations of interest: none. The datasets generated during and/or analyzed during the current study are available from the corresponding author or author of E.U. upon reasonable request. The datasets are also available from NCBI GEO data base. Disclosure Statement: The authors have nothing to disclose.

## ACKNOWLEDGEMENTS

This work was supported by Japan Society for the Promotion of Science (JSPS) Grant-in-Aid for Scientific Research (A, B and C) [grant numbers 17500533, 20300243, 24240096, 25282020], and Grant-in-Aid for Challenging Exploratory Research [grant number 21650196]. Founders had no role in study design, data collection and analysis, decision to publish, or preparation of the manuscript.

## SUPPLEMENTARY DATA

Supplementary Data are available.

## Notes

### Competing Interest Statement

The authors have declared no competing interest.

